# An efficient inducible model for the control of gene expression in renin cells

**DOI:** 10.1101/2024.03.26.585202

**Authors:** Silvia Medrano, Manako Yamaguchi, Hiroki Yamaguchi, Lucas Ferreira de Almeida, Curt D. Sigmund, Maria Luisa S. Sequeira-Lopez, R. Ariel Gomez

**Affiliations:** Child Health Research Center Department of Pediatrics, University of Virginia, Charlottesville, VA, USA; Department of Biology, University of Virginia, Charlottesville, VA, USA; Department of Physiology, Medical College of Wisconsin, Milwaukee, USA

**Keywords:** gene targeting, renin cells, renin deletion, arteriolar hypertrophy, lineage tracing

## Abstract

**Background:** Fate mapping and genetic manipulation of renin cells have relied on either non-inducible *Cre* lines that can introduce developmental effects of gene deletion or BAC transgene-based inducible models that may be prone to spurious and/or ectopic gene expression.

**Methods:** We generated an inducible mouse model in which *CreERT2* is under the control of the endogenous *Akr1b7* gene, an independent marker of renin cells.

**Results:** We evaluated the pattern of *Cre* expression in *Akr1b7^CreERT2/+^*;*R26R^mTmG/+^* mice in which Akr1b7^+^/renin^+^ cells become GFP+ upon tamoxifen administration. At E18.5 and P5, GFP was found in Juxtaglomerular cells, along the arterioles, and in the glomerular mesangium. In adult kidneys, GFP was present mainly in Juxtaglomerular cells. In mice treated with captopril and a low sodium diet to induce recruitment of renin cells, GFP extended along the afferent arterioles and in the mesangium. In addition, we deleted renin in adult mice and found a marked reduction in kidney renin expression and mean arterial pressure in mutant animals. When subjected to a homeostatic threat, mutant mice were unable to recruit renin^+^ cells. Most importantly, mice with renin deletion induced in the adult developed concentric vascular hypertrophy ruling out potential developmental effects on the vasculature due to the lack of renin.

**Conclusions:** Renin can be efficiently and specifically deleted in renin cells in adult life using our conditional model. *Akr1b7^CreERT2^* mice constitute an excellent model for the fate mapping of renin cells and for the spatial and temporal control of gene expression in renin cells.

## INTRODUCTION

Renin cells are essential for survival. In the adult unstressed mammal, renin cells are located in the afferent arterioles near the glomeruli, thus their name juxtaglomerular (JG) cells. JG cells synthesize and release renin, an enzyme-hormone crucial for the regulation of blood pressure and fluid electrolyte homeostasis (1). In addition, renin cells have important roles in tissue morphogenesis, kidney vascular development, tissue repair and regeneration, and innate immune defense (2, 3, 4, 5). Renin cells exhibit a high degree of plasticity. In response to homeostatic stress, cells of the renin lineage, such as vascular smooth muscle cells, mesangial cells, and pericytes, re-express renin, in a process known as recruitment (6). This response is usually sufficient to restore homeostasis. However, under conditions of chronic and persistent activation of the renin cell program, such as the permanent inhibition of the Renin-Angiotensin System (RAS), renin cells contribute to the development of concentric vascular hypertrophy (7, 8, 9, 10, 11, 12, 13, 14, 15). The genetic and epigenetic regulatory mechanisms that control the remarkable plasticity of renin cells are beginning to be unraveled although the involved signals need to be identified. To understand the plasticity of renin cells *in vivo* it is necessary to have a model that allows the temporal and spatial control of gene regulatory networks without developmental effects that may occur using constitutive *Cre* lines.

Gene targeting in renin cells has relied either on non-inducible Cre recombinase or bacterial artificial chromosome (BAC) transgene-based inducible mouse lines (6, 14, 16). In particular, the *Ren1^d Cre^* line developed in our lab (6), has been extensively used to delete specific genes in renin lineage cells, and to label cells with fluorescent markers for lineage tracing and cell isolation, providing significant information on renin cell identity and function (17, 18). However, non-inducible *Cre* recombinase mouse models can introduce confounding developmental effects of gene deletion in the adult animal, and transgenic lines may be prone to spurious and/or ectopic gene expression. To circumvent these problems, an efficient, specific, temporally and spatially conditional model is necessary.

Temporal control of gene expression can be regulated using a modified *Cre* recombinase fused to a mutated ligand-binding domain of the estrogen receptor (*CreERT2*) (19). This modified domain binds with high affinity the synthetic ER ligand tamoxifen, but not the endogenous estrogen. Upon tamoxifen administration, *Cre* is translocated into the nucleus and catalyzes the recombination at loxP-flanked alleles (19). For spatial control of gene expression, the *CreERT2* cassette is placed under the control of a cell-specific gene. The aldo-keto-reductase Akr1b7 is an independent marker of renin cells that is co-expressed with renin under different developmental, physiological, and pathological conditions (20, 21). It has been suggested that the function of Akr1b7 is to remove harmful aldehydes resulting from the high synthetic activity of renin cells (20, 21). During embryonic life, Akr1b7 and renin are expressed in the big intrarenal arteries, along arterioles, and in JG and mesangial cells, whereas in the adult both genes become confined to the JG area (20, 21). In addition, under physiological stress Akr1b7 is co-expressed with renin when renin cell descendants-smooth muscle cells, mesangial cells and pericytes-re-acquire the renin phenotype (21). Based on this information, we hypothesized that Akr1b7 was an excellent candidate for fate tracking and the temporal and spatial control of genes involved in the development and regulation of renin cells. Here, we report the characterization of a novel *Cre* mouse model, *Akr1b7^CreERT2^*, for the spatial and temporal regulation of gene expression in renin cells.

## METHODS

### Generation of Akr1b7^CreERT2^ mice

*Akr1b7^CreERT2^* mice were generated by inserting a *P2A-CreERT2* knock-in cassette immediately upstream of the TGA stop codon of the mouse *Akr1b7* gene followed by the endogenous 3’UTR using a homologous recombination-based technique (Ingenious Targeting Laboratory, Ronkonkoma, NY; Fig. 1A). Briefly, a targeting vector was constructed by subcloning a 9.1 kb region from an *Akr1b7* positively identified C57BL/6 fosmid clone (WI1-1577M19) into the 2.45 kb-iTL cloning vector derived from the pSP72 vector (Promega; Madison, WI). The region was designed so the long homology arm (LA) extended 4.9 kb upstream of the *P2A-CreERT2* knock-in cassette and the short homology arm (SA) extended 1.9 kb 3’ to a *LoxP-FRT*-flanked Neomycin cassette that was inserted downstream of the 3’UTR. The entire *P2A-CreERT2* knock-in cassette was confirmed by sequencing. Ten micrograms of the 15.8 kb targeting vector were linearized with *Not*I and transfected by electroporation into FLP C57BL/6 (BF1) embryonic stem cells for homologous recombination. After selection with G418 antibiotic, surviving clones were expanded for PCR analysis to identify recombinant ES clones. We confirmed the correct integration of the *Akr1b7^CreERT2^* knock-in allele by PCR analysis of expanded ES clones using primers: NEOGT: 5’-GTC CGT GTC GCGA AGT TCC TAT ACT TTC-3’ and A1: 5’-ACT AGT CAC TTT CTC ATG CAA GGG G-3’. Product size: 2.04 kb (Fig. 1A and B). The *Neo* cassette in the targeting vector was removed during ES clone expansion. We confirmed the retention of the *P2A-CreERT2* cassette in the ES clones by PCR using primers SQ1: 5’-TGA TAG CTC GCA GGC CTT CCT TC-3’ and FN2A: 5’-AAC TTC GCG ACA CGG ACA CAA TCC –3’. Product size: 2.61 kb (Fig.1A and B). We determined the copy number of the targeting vector in ES clones by Real-time PCR to confirm a single integration (Fig. 1C). Probe A was designed at the knock-in insertion site and therefore annealed only to the WT allele. One copy indicated that the clone was correctly targeted. Probe B was designed in the region of the long homology arm and annealed to both WT and targeted alleles. Clones with two copies were considered as having a single integration. Targeted iTL BF1 (C57BL/6 FLP) embryonic stem cells were microinjected into Balb/c blastocysts. Resulting chimeras with a high percentage black coat color were mated to C57BL/6N WT mice to generate germline *Neo* deleted mice. *Akr1b7^CreERT2^* mice were crossed with C57BL/6J mice or reporter mice on a C57L/6J background for characterization.

**Figure 1.**
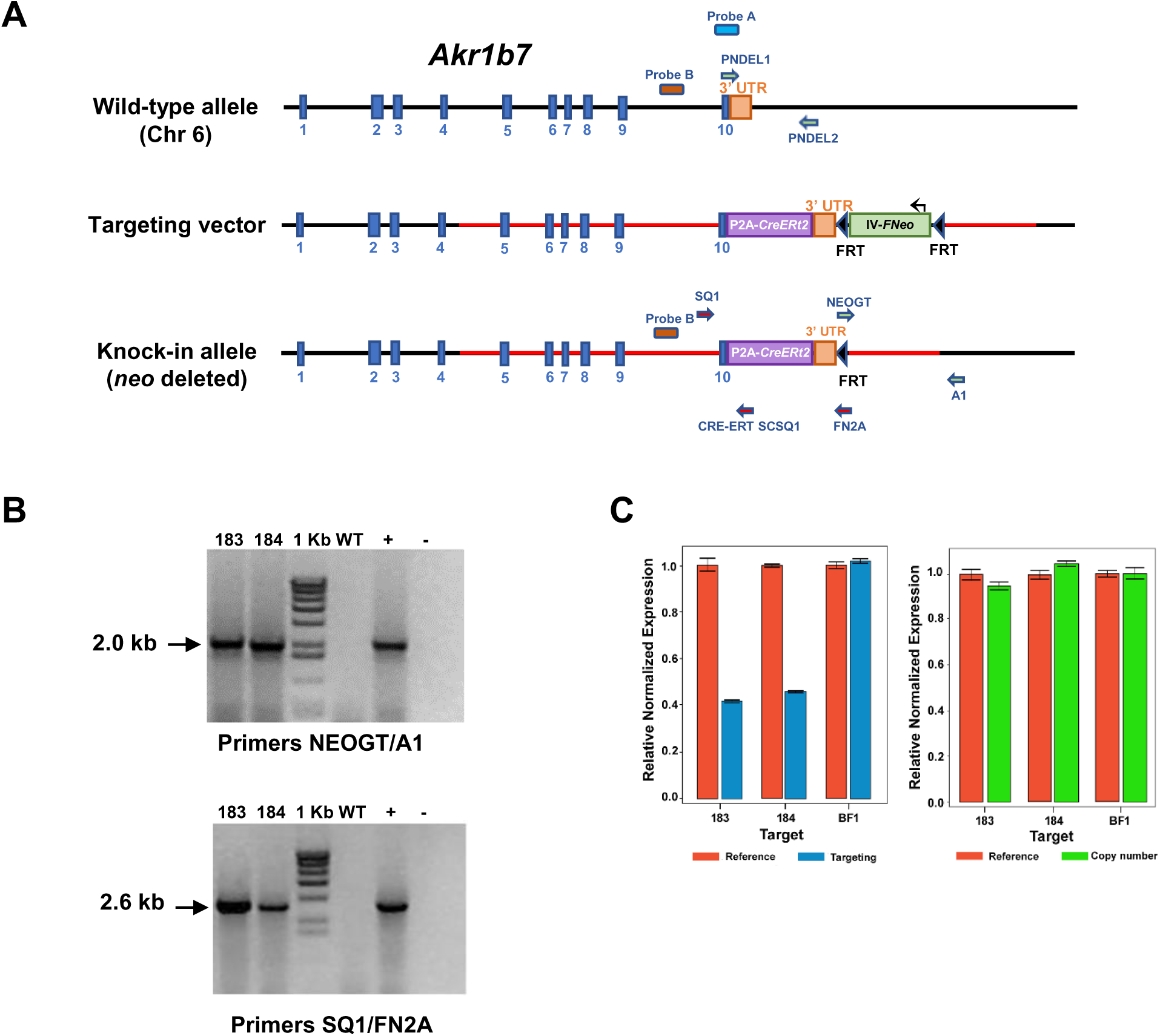
Generation of *Akr1b7^CreERT2^* mice. **A**: Schematic diagram of the targeting strategy for the generation of the *Akr1b7^CreERT2^* knock-in allele. **Top**. Map of the mouse *Akr1b7* gene in chromosome 6 showing the intron/exon structure. Exons are indicated by numbered blue boxes. **Middle.** For the generation of the targeting vector, a 9.1 kb region was first subcloned from a positively identified C57BL/6 fosmid clone (WI1-1577M19). Homology arms are indicated by red bars. The long homology arm extended 4.9 kb upstream of the *P2A-CreERT2* knock-in cassette and the short homology arm extended 1.9 kb 3’ to the *Neo* cassette. The *P2A-CreERT2* knock-in cassette was inserted immediately upstream of the TGA stop codon of the *Akr1b7* gene, followed by the endogenous 3’UTR. An *FRT*-flanked *Neo* selection cassette was inserted downstream of the 3’UTR. **Bottom**. Knock-in allele after *FRT*-directed *Neo* deletion in embryonic stem (ES) cells. **B. Top panel.** PCR analysis of two expanded ES clones (183 and 184) confirmed correct integration of the *Akr1b7^CreERT2^* knock-in allele. Wild-type (WT) DNA was used as a negative control. DNA from an individual clone (before expansion) was used as a positive control (+). No DNA was used as blank (-). Primers: NEOGT/A1. Product size: 2.0 Kb. **Bottom panel**. PCR analysis of expanded ES clones confirmed retention of the *P2A-CreERT2* cassette after expansion of the ES clones. Primers: SQ1/FN2A. Product size: 2.6 Kb. **C.** Analysis of copy number of targeting vector in ES clones by Real-time PCR. Probe A was designed at the knock-in insertion site and therefore annealed only to the WT allele. One copy indicated that the clone was correctly targeted. The WT sample, indicated as BF1, had 2 copies. Probe B was designed in the region of the long homology arm and annealed to both WT and targeted alleles. Clones with two copies were considered as having a single integration. The WT sample, indicated as BF1, had 2 copies. Clones 183, and 184 were correctly targeted and carried a single copy of targeting vector sequence.

### Genotyping of Akr1b7^CreERT2^ mice

Genotyping of the mice was conducted by PCR of DNA from tail biopsies using primers SQ1: 5’-TGA TAG CTC GCA GGC CTT CCT TC / CRE-ERT SCSQ1: 5’-ATC TTC AGG TTC TGC GGG AAA CCA-3’ to detect the *Akr1b7^CreERT2^* allele (product size 538 bp) and PNDEL1: 5’-CAG AGA AAC GAA GGG CAA CG-3’ / PNDEL2: 5’-GCT CCC TGA TGC CTC CTC TA-3’ (product size 838 bp) to detect the wild-type allele (Fig. 1A).

### Animals

To visualize the expression of *Akr1b7^CreERT2^* using fluorescent reporter genes, we generated 1) *Akr1b7^CreERT2/+^;R26R^mTmG/+^* animals by crossing *Akr1b7^CreERT2^* mice with the *R26R^mTmG^* Cre reporter mouse, strain # 007676, The Jackson Laboratory, Bar Harbor, ME (22); 2) *Akr1b7^CreERT2/+^;R26R^tdTomato/+^* animals by crossing *Akr1b7^CreERT2^* mice with the *R26R^tdTomato^* Cre reporter (Strain # 007909, also known as Ai9, The Jackson Laboratory); 3) *Akr1b7^CreERT2/+^;Ren1^cYFP^;R26R^tdTomato/+^* animals by crossing *Akr1b7^CreERT2^* mice with *Ren1^cYFP^;R26R^tdTomato^*reporter mice. In this line, the *Ren1^cYFP^* transgene labels cells actively expressing renin as we have previously shown (23). To delete renin in adult animals and simultaneously label renin cells with GFP, we generated *Akr1b7^CreERT2/+^;Ren1^cFl/−^;R26R^mTmG/+^* (Renin cKO*^GFP^*) by crossing *Akr1b7^CreERT2/CreERT2^*; *Ren1^c+/−^* to *Ren1^cFl/Fl (Homo)^*; *R26R^mTmG/mTmG^*. The *Ren1^c^ null* (*Ren1^c−/−^*) mouse line was generated by Takahashi et al. (9). The *Ren1^c^* floxed mouse were generated by Xu et al. (24). This crossing strategy yielded *Akr1b7^CreERT2/+^;Ren1^cFl/−^;R26R^mTmG/+^* mutant mice and *Akr1b7^CreERT2/+^;Ren1^cFl/+^;R26R^mTmG/+^* littermate controls. All animals used in this study were maintained in the C57BL/6 background.

All animals were housed in the University of Virginia’s vivarium facilities equipped with controlled temperature and humidity conditions in a 12 hours dark/light cycle. All procedures were performed per the Guidelines for the Care and Use of Laboratory Animals published by the United States National Institutes of Health (https://grants.nih.gov/grants/olaw/guide-for-the-care-and-use-of-laboratory-animals.pdf) and approved by the University of Virginia Animal Care and Use Committee.

### Blood Pressure Measurement

Mean arterial, systolic and diastolic blood pressure were measured in isoflurane anesthetized animals over a 10-min period from a catheter inserted into the right carotid artery using a RX104A transducer coupled to a data acquisition system and AcqKnowledge software (Biopac Systems, Inc., Goleta, CA).

### Blood analysis

Animals were anesthetized with tribromoethanol (300 mg/kg). Blood was collected by cardiac puncture and placed into heparinized plasma separator tubes (BD Microtainer, Becton Dickinson, Franklin Lakes, NJ). For mice subjected to blood pressure analysis, blood was collected from the right carotid artery under isoflurane anesthesia. Tests for basic metabolic panel were performed by the University of Virginia Hospital clinical laboratory.

### Plasma renin

The plasma renin concentration was quantified using an ELISA kit for mouse renin 1 (Ray Biotech, Norcross, GA) as previously described (15).

### Histological and immunohistochemical analyses

Mice were anesthetized with tribromoethanol (300 mg/kg). Kidneys were removed, fixed overnight in Bouin’s solution at room temperature (RT) or 4% paraformaldehyde (PFA) at 4°C, and embedded in paraffin. Sections (5 µm) from Bouin’s-fixed, paraffin-embedded kidneys were stained with Hematoxylin and Eosin (MilliporeSigma, Burlington, MA) to examine kidney morphology. Immunostaining was performed on 5 µm sections of Bouin’s-fixed, paraffin-embedded kidneys using a rabbit polyclonal anti-mouse renin antibody (1:500) generated in our laboratory (25), or mouse anti α-SMA monoclonal antibody (1:10,000; MilliporeSigma), and biotinylated secondary goat anti–rabbit IgG or horse anti–mouse IgG (1:200; Vector Laboratories, Newark, CA) for renin or α-SMA respectively. Staining was amplified using the Vectastain ABC kit (Vector Laboratories) and developed with 3,3-diaminobenzidine (MilliporeSigma). The sections were counterstained with hematoxylin (MilliporeSigma), dehydrated, and mounted with Cytoseal XYL (Thermo Fisher Scientific, Waltham, MA).

GFP expression after *Cre* recombination was visualized in frozen sections. Tissues were fixed in 4% PFA for 1 hour at 4°C. After washing, the samples were incubated in 30% sucrose overnight at 4°C and frozen in O.C.T. (Thermo Fisher Scientific, Waltham, MA). The frozen blocks were sectioned at 12 μm thickness and mounted in phosphate-buffered saline (PBS).

### In situ hybridization

To generate the probe for *Akr1b7 in situ* hybridization, we synthesized a 450 bp DNA fragment by PCR using cDNA from wild-type C57BL/6 mouse kidneys and primers AATTAACCCTCACTAAAGGGTGACCAACCAGATTGAGAGC and TAATACGACTCACTATAGGGCAGTATTCCTCGTGGAAAGGAT containing a 3′ T3 promoter a 5′ T7 promoter sequences respectively. Digoxigenin (DIG)-labeled RNA sense and antisense probes were generated by *in vitro* transcription using DIG RNA Labeling Mix and T3 or T7 Polymerases (MilliporeSigma). *In situ* hybridization was performed as previously described (15). Briefly, 7 μm 4% PFA-fixed, paraffin-embedded kidney sections were deparaffinized, rehydrated and postfixed with 4% PFA, followed by acetylation (0.375% acetic anhydride) and permeabilization with proteinase K (10 μg/mL). After preincubation with hybridization buffer (500 ng/mL in hybridization buffer of 50% formamide, 5× SSC, 50 μg/mL yeast transfer RNA, 1% SDS, 50 μg/mL heparin), sections were incubated with the DIG-labeled sense or antisense riboprobes at 55°C overnight. The sections were washed and then incubated with anti–digoxigenin-alkaline phosphatase antibody (1:4,000, MilliporeSigma) overnight at 4°C. After washing, sections were treated with levamisole and incubated with BM Purple (MilliporeSigma) until the signals were visible. Sections were fixed with 0.2% glutaraldehyde ^+^ 4% PFA and mounted with Glycergel Mounting Medium (Agilent Technologies, Santa Clara, CA). The antisense probe generated with T7 polymerase showed the specific signal. The sense probe generated with T3 polymerase showed no signal.

### Microscopy

Tissue sections were visualized using a Zeiss Imager M2 microscope equipped with the AxioCam 305 color and AxioCam 506 mono cameras (Zeiss, Oberkochen, Germany).

### Quantitative assessment

We determined the juxtaglomerular area index (JGAi) as the number of YFP or tdTomato positive JG areas ÷ total number of glomeruli × 100 in 12 randomly selected 20x cortex images for each animal under basal physiological conditions. To quantify the renin immunostaining in kidney sections, we used the magic wand tool of Photoshop software to automatically select positive areas in 12 randomly selected 20x cortex images. The measurements were normalized for the total cortex area in each image and expressed as percentages. All the image analysis parameters were kept constant among different samples.

### Tissue-clearing and whole-mount immunostaining

For the tissue-clearing and whole-mount immunostaining of *Akr1b7^CreERt2^;R26R^tdTomato^* mouse kidneys, we used the updated clear, unobstructed brain/body imaging cocktails and computational analysis (CUBIC) protocol (26). The mice were anesthetized with tribromoethanol (300 mg/kg) and perfused with 20 mL of PBS and 30 mL of 4% PFA via the left ventricle of the heart. Kidneys were removed, divided into two sections each, and fixed overnight in 4% PFA. All subsequent steps were performed with gentle shaking. The samples were washed with PBS for 6 hours followed by immersion in CUBIC-L (10 wt% *N*-butyldiethanolamine and 10 wt% Triton X-100) at 45°C for 5 days. After the samples were washed with PBS for 6 hours, they were placed in blocking buffer (PBS, 1.0% bovine serum albumin, and 0.01% sodium azide) overnight at RT. Then the samples were immersed in immunostaining buffer (PBS, 0.5% Triton X-100, 0.25% bovine serum albumin, and 0.01% sodium azide) containing 1:100 diluted Anti-Actin, α-Smooth Muscle (Acta2) - Fluorescein (FITC) antibody (F3777, Sigma-Aldrich) for 6 days at RT. After washing again with PBS for 6 hours, the samples were immersed in 1:1 diluted CUBIC-R+ [T3741 (45 wt% 2,3-dimethyl-1-phenyl-5-pryrazolone, 30 wt% nicotinamide and 5 wt% *N*-butyldiethanolamine), Tokyo Chemical Industry, Tokyo, Japan] overnight at RT. The samples were then immersed in CUBIC-R+ at RT for 2 days.

Macroscopic whole-mount images were acquired with a light-sheet fluorescence (LSF) microscope (ZEISS Lightsheet7, Zeiss). The samples imaged in mounting solution (RI:1.520) (M3294, Tokyo Chemical Industry) with Clr Plan-Neofluar 20x/1.0 Corr nd=1.53 detection optics. The voxel resolution was as follows: x = 0.656 µm, y = 0.656 µm, z = 1.0 µm (zoom × 0.36); x = 0.169 µm, y = 0.169 µm, z = 0.560 µm (zoom × 1.4, 2 x 2 tiles). The FITC signals of α-smooth muscle actin expressing cells were measured by excitation with 488 nm lasers. The TdTomato signals were measured by excitation with 561 nm lasers.

Three-dimensionally rendered images were visualized, captured with Imaris software (Version 10.0.0, Bitplane, Belfast, United Kingdom). Raw image files obtained from Zeiss LSF microscope (.czi) were converted into Imaris files (.ims) using Imaris File Converter 10.0.0. Tilescan images were stitched using Imaris Stitcher 10.0.0. The image processing by Imaris software was performed as previously described (27). The reconstituted 3D images were cropped to a region of interest using the crop function. The snapshot and animation functions were used to capture images and videos, respectively.

### RNA isolation and Real-time RT-PCR analysis

Quantitative Real Time PCR (qPCR) analysis was performed in samples from kidney cortices. Total RNA was extracted using TRIzol reagent (Thermo Fisher Scientific) and RNeasy Mini Kit (Qiagen, Germantown MD). Reverse transcription was performed using oligo(dT) primers and M-MLV Reverse Transcriptase (Promega, Madison, WI) at 42°C for 1 hour according to the manufacturer’s instructions. qPCR was performed using SYBR Green I (Thermo Fisher Scientific) in a CFX Connect system (Bio-Rad Laboratories, Hercules, CA). PCR was performed using the following primers; *Ren1*, forward: 5′-ACAGTATCCCAACAGGAGAGACAAG-3′, reverse: 5′-GCACCCAGGACCCAGACA-3′; *Rps14*, forward: 5′-CAGGACCAAGACCCCTGGA-3′, reverse: 5′-ATCTTCATCCCAGAGCGAGC-3′. *Ren1* mRNA expression was normalized to *Rps14* expression, and the changes in expression were determined by the ΔΔCt method and reported as relative expression compared with control mice.

### Statistical Analysis

Data are presented as means ± SD. Statistical analysis was performed using GraphPad Prism 10 software (GraphPad Software, San Diego, CA). Comparisons between two groups were performed by two-tailed unpaired Student’s *t*-test. Comparisons between more than two groups were performed by Brown-Forsythe and Welch ANOVA followed by Dunnett’s T3 multiple comparisons test. *P* values < 0.05 were considered statistically significant.

## RESULTS

### Akr1b7^CreERT2^ mice exhibit normal kidney structure and the same expression levels of Akr1b7 than wild-type mice

To achieve temporal control of gene expression in renin cells, we generated *Akr1b7^CreERT2^* mice by inserting a *P2A-CreERT2* knock-in cassette immediately upstream of the TGA stop codon of the mouse *Akr1b7* gene followed by the endogenous 3’UTR using a homologous recombination approach. Figure 1A shows a schematic diagram of the of the mouse *Akr1b7* gene, the targeting vector and the final knock-in allele. We confirmed the correct insertion and retention of the *Akr1b7^CreERT2^* knock-in allele by PCR in embryonic stem (ES) cells (Fig. 1B) and the presence of a single insertion of the targeted allele (Fig. 1C). Homozygous and heterozygous *Akr1b7^CreERT2^* mice were viable, fertile, normal in size, and did not display any gross physical or behavioral abnormalities. In addition, the kidneys of *Akr1b7^CreERT2/+^* animals exhibited normal size and morphology, and same levels of expression of endogenous *Akr1b7* than wild-type mice (Suppl. Fig. 1).

### Akr1b7^CreERT2^ mice display specific Cre recombinase activity in renin cells

We first looked at the pattern of Cre expression during development and in the adult using *Akr1b7^CreERT2^;R26R^mTmG^* reporter mice where *Akr1b7* expressing cells become labeled with GFP upon *Cre* recombinase activation with tamoxifen and all non-recombined cells express tdTomato. As mentioned above, expression of Akr1b7 in the kidney follows that of renin throughout development and adulthood, and when newly transformed cells acquire the renin phenotype in response to a physiological threat (20, 21). Figure 2 shows the GFP and tdTomato expression in frozen kidney sections during postnatal life (P5 to P60). Neonates received three intragastric injections of 50 µg tamoxifen at P1, P2 and P3, and were analyzed at P5. We observed GFP signal along the arterioles, in the glomerular mesangium and in JG cells (Fig. 2A). Adult mice (P30–P60) were injected with tamoxifen (2mg/20g body weight) for 3 consecutive days and analyzed 2 days after the last injection. At P30, GFP signal was mainly confined to JG cells and some arterioles (Fig. 2B). Mice injected with corn oil (vehicle) did not show any GFP signal, indicating that *Cre* recombination did not occur in the absence of tamoxifen (Fig. 2C). Next, we examined the pattern of *Cre* expression in conditions of physiological stress. Sixty-day-old mice were treated with 0.5 mg/mL of the angiotensin-converting enzyme inhibitor captopril (Sigma-Aldrich, St. Louis, MO) in the drinking water and fed a low-NaCl (0.1%) diet for 8 days to induce recruitment of renin cells, and given tamoxifen i.p. on days 3, 4 and 5 of recruitment. As expected for recruited animals, we observed high GFP expression in JG cells, along the afferent arterioles and the intraglomerular mesangium (Fig. 2D).

**Figure 2.**
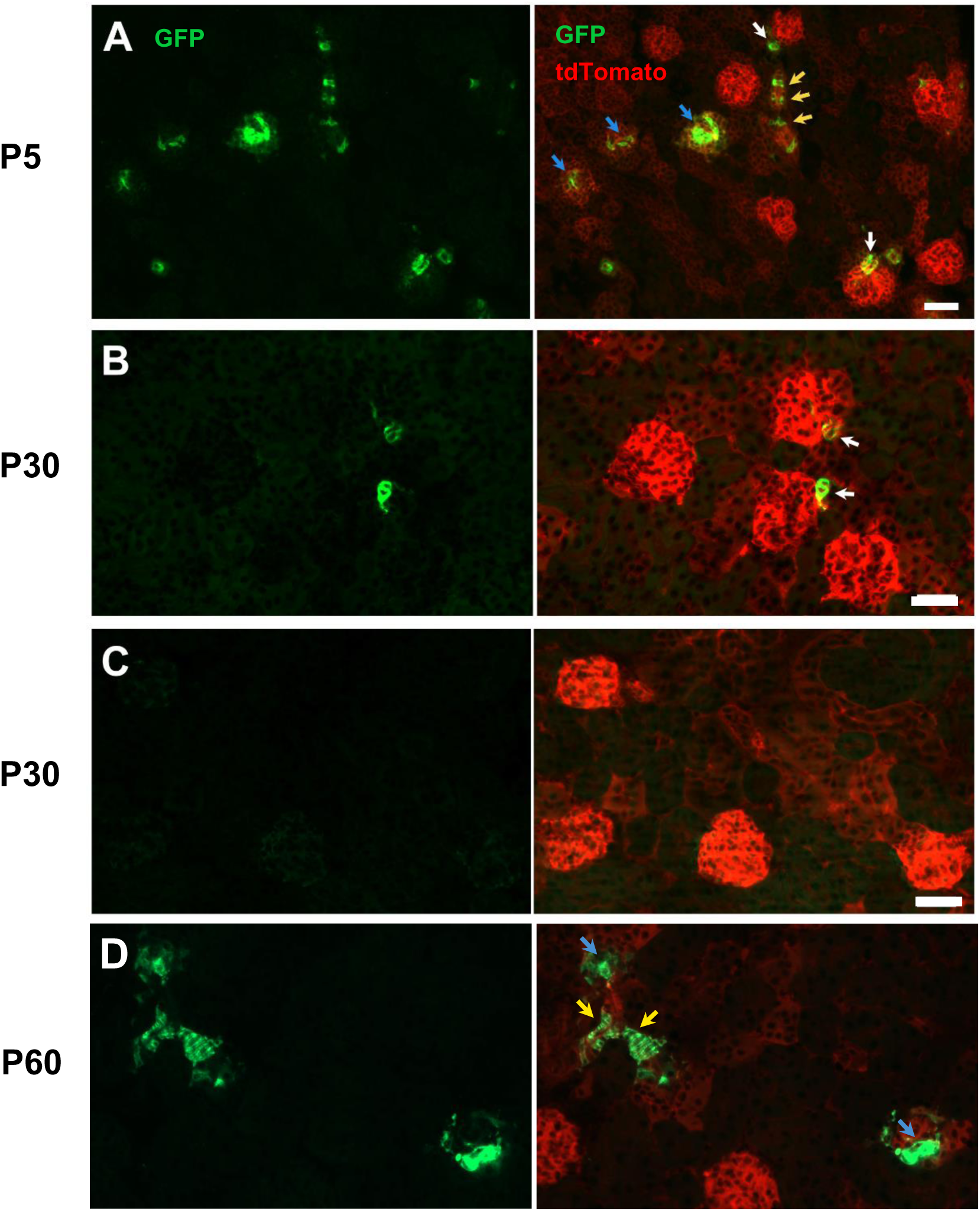
Pattern of Cre expression during postnatal life in *Akr1b7^CreERT2^* reporter mice. GFP and tdTomato expression in frozen sections of kidneys from tamoxifen- (**A**, **B** and **D**) and corn oil (vehicle) (**C**) Akr1b7*^CreERT2^*;R26R*^mTmG^* treated mice. In this reporter mouse, *Akr1b7* expressing cells become labeled with GFP upon *Cre* recombinase activation with tamoxifen and non-recombined cells express tdTomato. Left panels show GFP+ cells and right panels show GFP+ recombinant cells and tdTomato+ non-recombinant cells. **A.** At P5, we observed GFP signal along the vasculature (yellow arrows), in the intraglomerular mesangium (blue arrows) and juxtaglomerular (JG) areas (white arrows). Bar: 75 µm. **B.** At P30, GFP signal was mainly confined to JG areas (white arrows) and some arterioles (yellow arrows). Bar: 50 µm. **C.** P30 kidney from a mouse injected with corn oil. No GFP signal was observed indicating that Cre recombination did not occur in the absence of tamoxifen. Bar: 50 µm. **D.** Two-month-old mouse treated with 0.5 mg/mL of the angiotensin-converting enzyme inhibitor captopril in the drinking water and fed a low-NaCl (0.1%) diet for 8 days to induce recruitment of renin cells. GFP expression extended along the afferent arterioles (yellow arrows) and the intraglomerular mesangium (blue arrows). Bar: 50 µm. These results indicate that in *Akr1b7^CreERT2^* mice, Cre is expressed in the same areas as the endogenous Akr1b7 under basal conditions and under homeostatic stress. In addition, Cre expression is tightly regulated by tamoxifen.

Akr1b7 is expressed in a few organs outside the kidney, most prominently in the cortex of the adrenal gland (28). In P30 *Akr1b7^CreERT2^;R26R^mTmG^* treated with tamoxifen as described for Figure 2B, we observed strong GFP expression in the cortex of adrenal glands where the signal was restricted to the zona fasciculata (Suppl. Fig. 2) as reported for the endogenous Akr1b7 protein.

To determine the spatial distribution of renin cells and their topological relationships with glomeruli, tubules, vasculature and interstitium, we performed tissue-clearing-based three-dimensional (3D) imaging using additional reporter mice. We generated *Akr1b7^CreERT2^;R26R^tdTomato^*mice where *Akr1b7* expressing cells are labeled with red fluorescence upon *Cre* activation by tamoxifen. We used the updated clear, unobstructed brain/body imaging cocktails and computational analysis (CUBIC) protocol (26, 27) and light-sheet fluorescence (LSF) microscopy to visualize combined reporter expression and immunofluorescence for Acta2 to label the vasculature. This technique allowed us to visualize intact kidneys without slicing the tissue. Figure 3 shows Imaris software (Bitplane) reconstructed high magnification 3D images of kidneys from mice under basal conditions and after administration of Captopril (0.5 mg/mL H_2_O) + low Na^+^ (0.1%) diet for 7 days to induce recruitment of renin cells. Under basal conditions, tdTomato signal was clearly localized to the tip of the arterioles in the JG areas after tamoxifen administration. Under physiological stress, tdTomato signal was much brighter, and the labelled JG areas looked larger compared to controls. In addition, the tdTomato signal extended along the arterioles as expected in recruited mice. Supplementary Video S1 shows the reconstructed 3D images at low magnification allowing the continuous observation of tomographic images of *Akr1b7^CreERt2^;R26R^tdTomato^* kidneys in all directions. In summary, 3D imaging experiments showed tdTomato signal where Akr1b7 is expected to be expressed in both control and recruited mice. In addition, this 3D imaging approach provided a comprehensive view of the kidney vasculature and a better visualization of the complex distribution of renin cells under basic conditions and physiological stress in the intact tissue.

**Figure 3.**
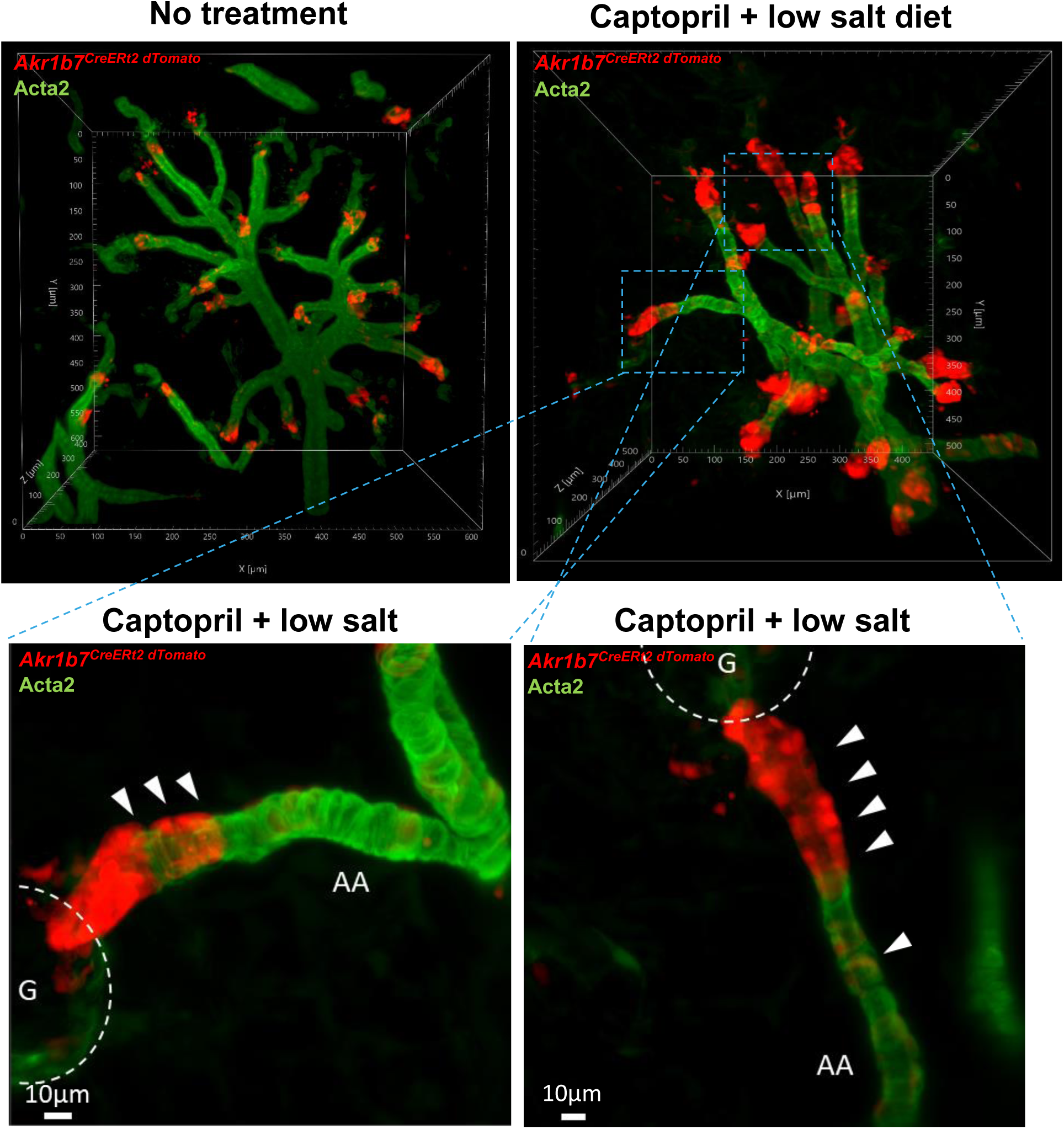
Tissue clearing and 3D imaging of kidneys from Akr1b7*^CreERT2^*;R26R*^tdTomato^* mice under normal conditions and physiological stress. We used the updated clear, unobstructed brain/body imaging cocktails and computational analysis (CUBIC) protocol and light-sheet fluorescence (LSF) microscopy to visualize combined reporter expression and immunofluorescence for Acta2 to label the vasculature. Shown are Imaris software (Bitplane) reconstructed high magnification 3D images of kidneys from three-month-old mice under basal conditions and after administration of captopril (0.5 mg/mL) + low Na^+^(0.1%) diet for 7 days to induce recruitment of renin cells. Mice received three consecutive i.p. injections of tamoxifen (2mg/20g BW) on days 1, 2 and 3 of the experiment. Red fluorescence indicates Akr1b7*^CreERT2^* expression upon tamoxifen administration and FITC shows immunofluorescence for Acta2. Under basal conditions, tdTomato signal was clearly localized to the tip of the arterioles in the JG areas. Under physiological stress, tdTomato signal was much brighter, and the labelled JG areas looked larger compared to controls. In addition, the tdTomato signal extended into the arterioles as expected in recruited mice (white arrowheads). See also Supplementary Videos 1 and 2. AA; afferent arteriole, G; glomerulus.

Next, we investigated the pattern of *Cre* expression during embryonic life using *Akr1b7^CreERt2^;R26R^mTmG^* reporter mice. Female mice received two consecutive i.p. injections of 2 mg/40g BW tamoxifen on days 15.5 and 16.5 of pregnancy, and embryos were studied at E18.5. Figure 4 shows images of frozen sections of kidneys and adrenal glands. Within the kidney, we observed GFP expression primarily along the vasculature, in the glomerular mesangium, and in JG areas of mature glomeruli. In addition, we found GFP expression in the zona fasciculata of the embryonic adrenal cortex. These results are consistent with the pattern of expression of Akr1b7 and renin at this stage of embryonic life.

**Figure 4.**
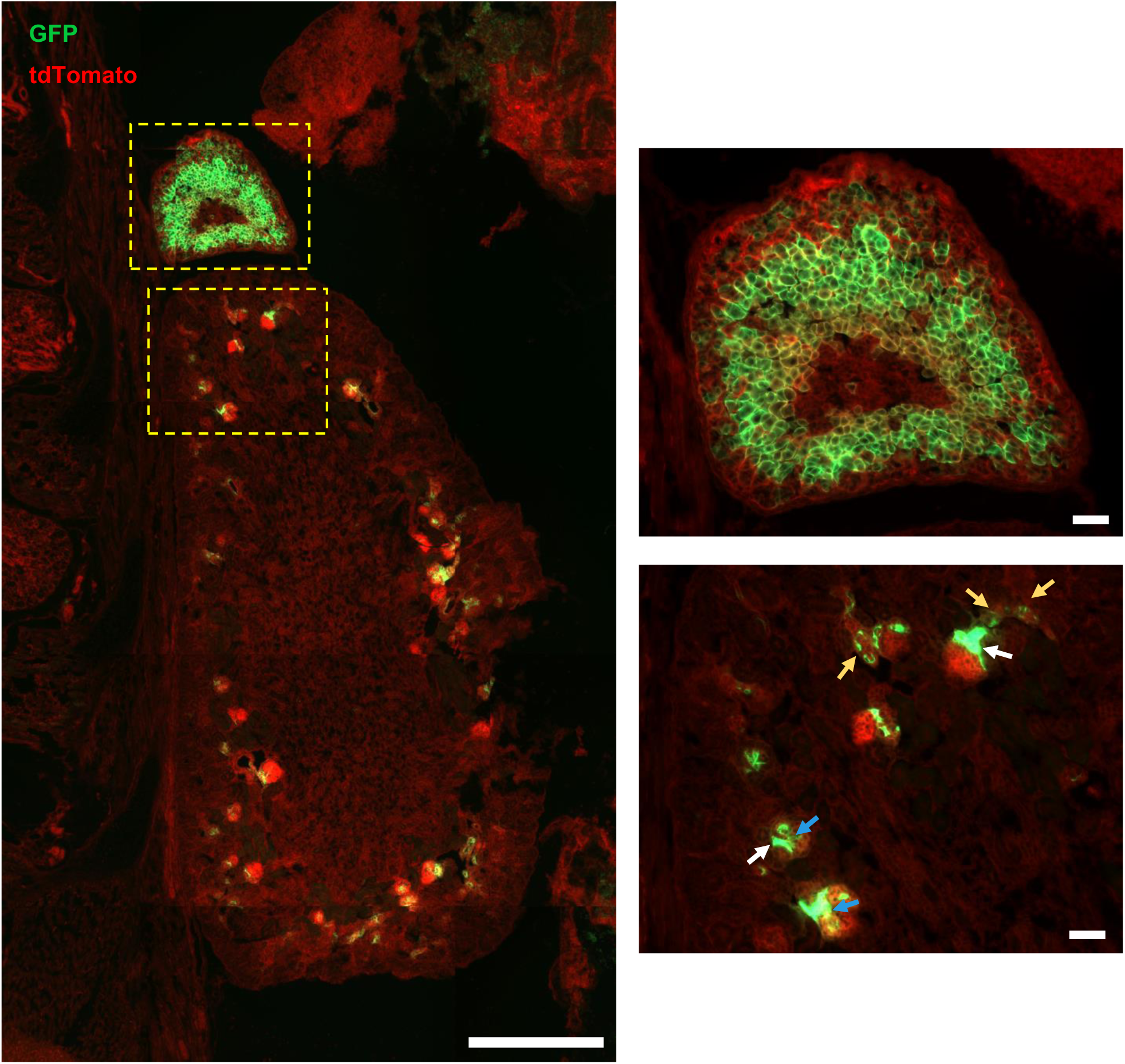
Pattern of Cre expression during embryonic life in *Akr1b7^CreERT2^* reporter mice. GFP and tdTomato expression in frozen sections of kidneys and adrenal glands from tamoxifen-treated *Akr1b7^CreERt2^;R26R^mTmG^* E18.5 embryos. Female mice received two consecutive i.p. injections of 2mg/40g BW tamoxifen on days 15.5 and 16.5 of pregnancy. **Left panel:** Low magnification image of the kidney and adrenal gland. Bar: 500 µm. **Right panels**: High magnification images of the areas indicated in the left panel. In the kidney, we observed GFP signal along the arteries and arterioles (yellow arrows), in the glomerular mesangium (blue arrows) and JG areas (white arrows) of mature glomeruli. In addition, GFP was expressed in the zona fasciculata of the embryonic adrenal cortex. Bars: 50 µm. A total of 5 E18.5 embryos were stutied. These results are consistent with the pattern of expression of endogenous Akr1b7 during embryonic life.

In all, our results indicate that, in *Akr1b7^CreERT2^* mice, *Cre* is expressed in the same areas reported for the endogenous Akr1b7 during development and in the adult both under basal conditions and under physiological stress. In addition, *Cre* activity is tightly regulated by tamoxifen.

### *Cre* and *Renin* highly co-localize in the kidney of *Akr1b7^CreERt2^* mice in the basal state and under physiological stress

Next, we asked whether reporters for *Akr1b7^CreERT2^* and renin co-localize in the kidney of these mice. To answer this question, we generated *Akr1b7^CreERT2^;Ren1^cYFP^;R26R^tdTomato^* reporter mice where cells actively expressing renin are labeled with YFP, as we have previously shown (23), and *Cre* expressing cells can be labeled with tdTomato upon tamoxifen administration. In these experiments, tamoxifen (2mg/20g BW) was administered i.p. on 3 consecutive days and mice were studied 3 days after the last injection. A group of mice were treated with captopril (0.5 mg/mL H_2_O) + low Na^+^(0.1%) diet for 8 days and tamoxifen was administered on days 3, 4 and 5 of recruitment. Figure 5 shows the images of frozen kidney sections from *Akr1b7^CreERT2^;Ren1^cYFP^;R26R^tdTomato^* mice under basal conditions and under physiological stress. We found that *Cre* (tdTomato) and Renin (YFP), highly localized to the same areas: under basal conditions in JG cells, and under physiological stress in JG cells, along the arterioles and the intraglomerular mesangium (Fig. 5A). The juxtaglomerular area index (JGAi), determined as the number of renin positive JG areas ÷ total number of glomeruli × 100, is an established method to quantitate and compare levels of renin expression in mice (29, 30). To quantify the degree of co-localization of *Cre* (tdTomato) and Renin (YFP) and gain insight into the rate of recombination in *Akr1b7^CreERT2^* mice, we compared the JGAi calculated using both reporters. We estimated the JGAis in 12 randomly selected 20x cortex images for each animal under basal physiological conditions. We found no significant differences between the JGAi for renin (YFP) positive cells and *Cre* (tdTomato) positive cells suggesting a high level of co-localization of the expression of the two genes (Fig. 5B). In addition, all YFP (renin) positive cells were positive for tdTomato suggesting that *Akr1b7^CreERT2^* mice exhibit a high rate of recombination.

**Figure 5.**
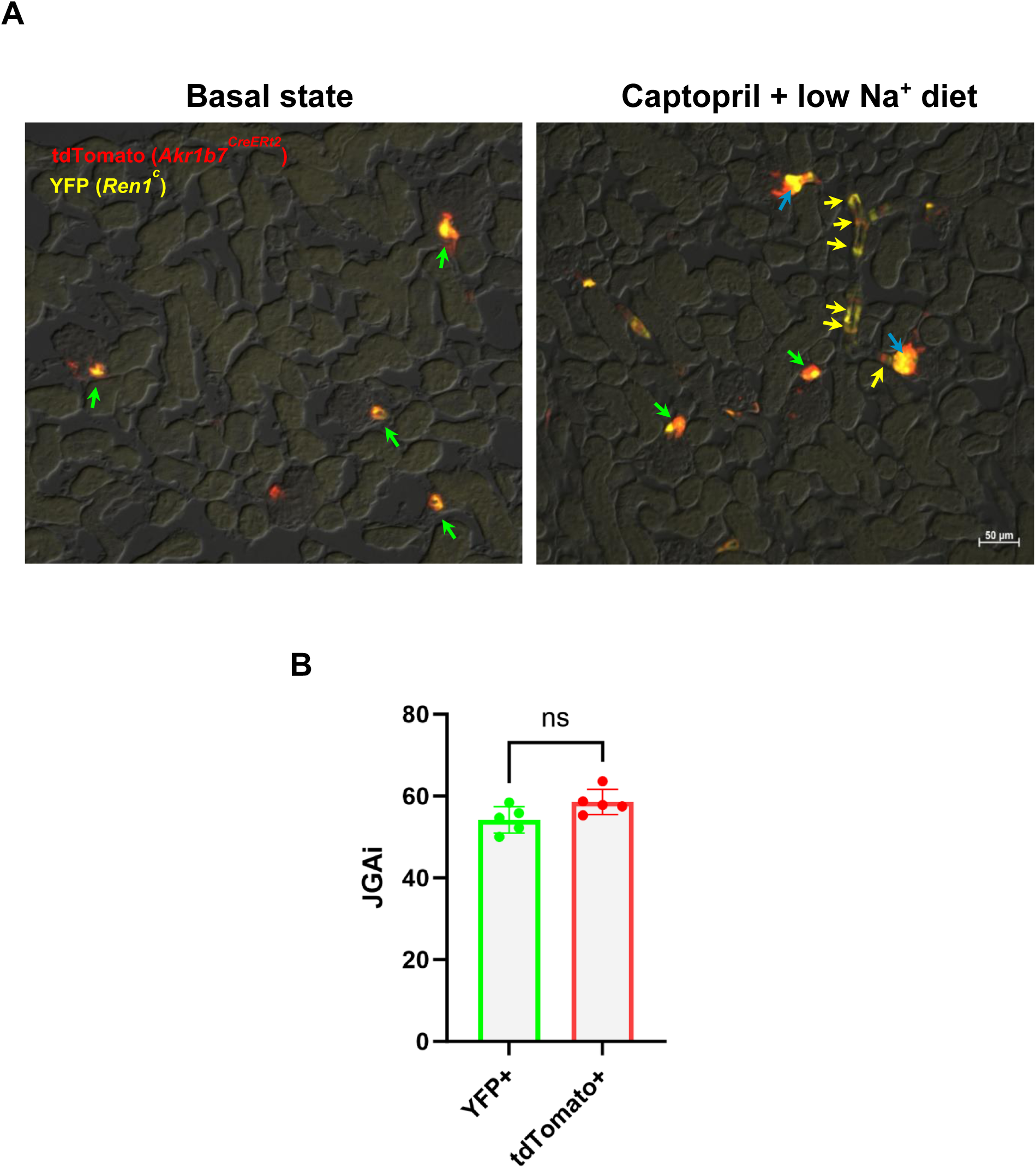
*Cre* and *Renin* highly co-localize in the kidney of *Akr1b7^CreERt2^* mice in the basal state and under physiological stress. **A.** YFP and tdTomato expression in frozen sections of kidneys from tamoxifen-treated *Akr1b7^CreERt2/+^;Ren1^cYFP^;R26R^tdTomato/+^* reporter mice. Cre (tdTomato) and Renin (YFP), highly localize to the same areas: under basal conditions in JG cells (green arrows), and under physiological stress in JG cells (green arrows), along the arterioles (yellow arrows) and the intraglomerular mesangium (blue arrows). All YFP (renin) positive cells were positive for tdTomato. These observations strongly suggest that in our model the reporters follow exactly the same pattern of expression as the endogenous Akr1b7 and renin. **B.** Juxtaglomerular area index (JGAi) calculated using YFP^+^ cells or tdTomato^+^ cells. We calculated the JGAis in 12 randomly selected 20x kidney cortex images for each animal under basal physiological conditions. We found no significant differences between the JGAi for renin (YFP) positive cells and Cre (tdTomato) positive cells. Umpaired Student’s *t*-test *P*>0.05; n=5; 3 females and 2 males; 34-52 days old.

These results clearly show that in *Akr1b7^CreERT2^;Ren1^cYFP^;R26R^tdTomato^* mice the reporters for *Cre* and renin are highly co-localized, confirming the renin cell specificity and high rate of recombination of our model.

### Renin can be efficiently deleted in the kidney of adult *Akr1b7^CreERT2^* mice

The effect of renin deletion has been extensively studied using mouse models of global renin deletion or constitutive Cre-Loxp systems (9, 13, 14, 15). One limitation in these studies is that renin is deleted throughout the life of the mouse, not only in the kidney but in other organs and tissues. This could result in confounding developmental effects when studying the response to renin deletion in the adult animal. To circumvent this problem, we used the *Akr1b7^CreERT2^* model to generate *Akr1b7^CreERT2^;Ren1^cFl/−^;R26^mTmG^* mice. In this mouse, the *Ren1^c^* floxed allele can be deleted and *Akr1b7^CreERT2^* expressing cells can be simultaneously labeled with GFP upon tamoxifen administration in a time controlled manner.

First, we conducted a short-term study to compare tamoxifen-treated *Akr1b7^CreERT2/+^;Ren1^cFl/^;R26^mTmG/+^* with *Akr1b7^CreERT2/+^;Ren1^c+/−^;R26^mTmG/+^* controls. Renin hemizygous mice are normal and therefore can be used as controls in this study (9). Adult mice received tamoxifen (2mg/20g BW) i.p. on 5 consecutive days and were analyzed after a washout period of 10–15 days. A group of mice were treated with captopril (0.5 mg/mL H_2_O) + low Na^+^ (0.1%) diet for the last 8 days of the experiment to induce recruitment of renin cells. Figure 6A shows immunostaining for Renin in kidney sections from control and renin mutant mice (Renin cKO*^GFP^*) under basal conditions and physiological stress. We observed a marked decrease in renin signal in kidneys of Renin cKO*^GFP^* mutant mice under both conditions compared to controls. The quantification of areas of Renin positive immunostaining is shown in Figure 6B. We observed a significant decrease in areas of renin immunostaining in mutant mice compared to control animals under basal conditions (Control-basal: 0.159±0.040%; n=8; Mutant-basal: 0.040±0.018; n=9; *P*<0.001). Similarly, renin immunostaining under recruitment conditions is significantly lower in Renin cKO*^GFP^* mutant mice compared to controls (Control-recruited: 0.441±0.139%; n=6; Mutant-recruited: 0.069±0.040; n=5; *P*<0.01). Figure 6B also shows that under physiological stress, control mice exhibited significantly higher levels of renin expression compared to basal conditions (Control-basal: 0.159±0.040%; n=8; Control-recruited: 0.441±0.139%; n=6; *P*<0.05). The renin levels in mutant recruited mice were not significantly different from those in non-recruited animals (Mutant-basal: 0.040±0.018; n=9; Mutant-recruited 0.069±0.040; n=5; *P*>0.05). In addition to the changes at the protein level, we observed a significant decrease in renin mRNA measured by qPCR in Renin cKO*^GFP^* mutant mice kidney cortices compared to controls in both the basal state (Control-basal: 0.92±0.26; n=8; Mutant-basal: 0.17±0.04; n=5; *P*<0.001) and under physiological threat (Control-recruited: 5.27±1.65; n=5; Mutant-recruited: 0.45±0.05; n=4; *P*<0.05) (Fig. 7A). As expected, we found that under physiological stress, control mice exhibited significantly higher levels of renin mRNA compared to basal conditions (Control-basal: 0.92±0.26; n=8; Control-recruited: 5.27±1.65; n=5; *P*<0.05). Renin mRNA levels in mutant recruited mice were significantly higher than in non-recruited animals (Mutant-basal: 0.17±0.04; n=5; Mutant-recruited: 0.45±0.05; n=4; *P*<0.001), suggesting that the animals are attempting to increase renin synthesis. In addition, we found that circulating renin levels followed a similar trend: Control-basal: 53,714±27,908; n=7 *vs* Mutant-basal: 12,466±7,035; n=9; *P*<0.05); Control-recruited: 309,954±122,023; n=5 *vs* Mutant-recruited: 44,375±12,955; n=5; *P*<0.05); Control-basal: 53,714±27908; n=7 *vs* Control-recruited: 309,954±122,023; n=5; *P*<0.05; Mutant-basal: 12,466±7,035; n=9 vs Mutant-recruited: 44,375±12,955; n=5; *P*<0.05 (Fig. 7B). In line with our renin mRNA and protein results, we found that renin mutant mice exhibited a significant decrease in arterial blood pressure under basal conditions compared to controls: Control-basal: 82.7±2.6; n=7 vs Mutant-basal: 68.9±5.3; n=9; *P*<0.005 (Fig. 7C).

**Figure 6.**
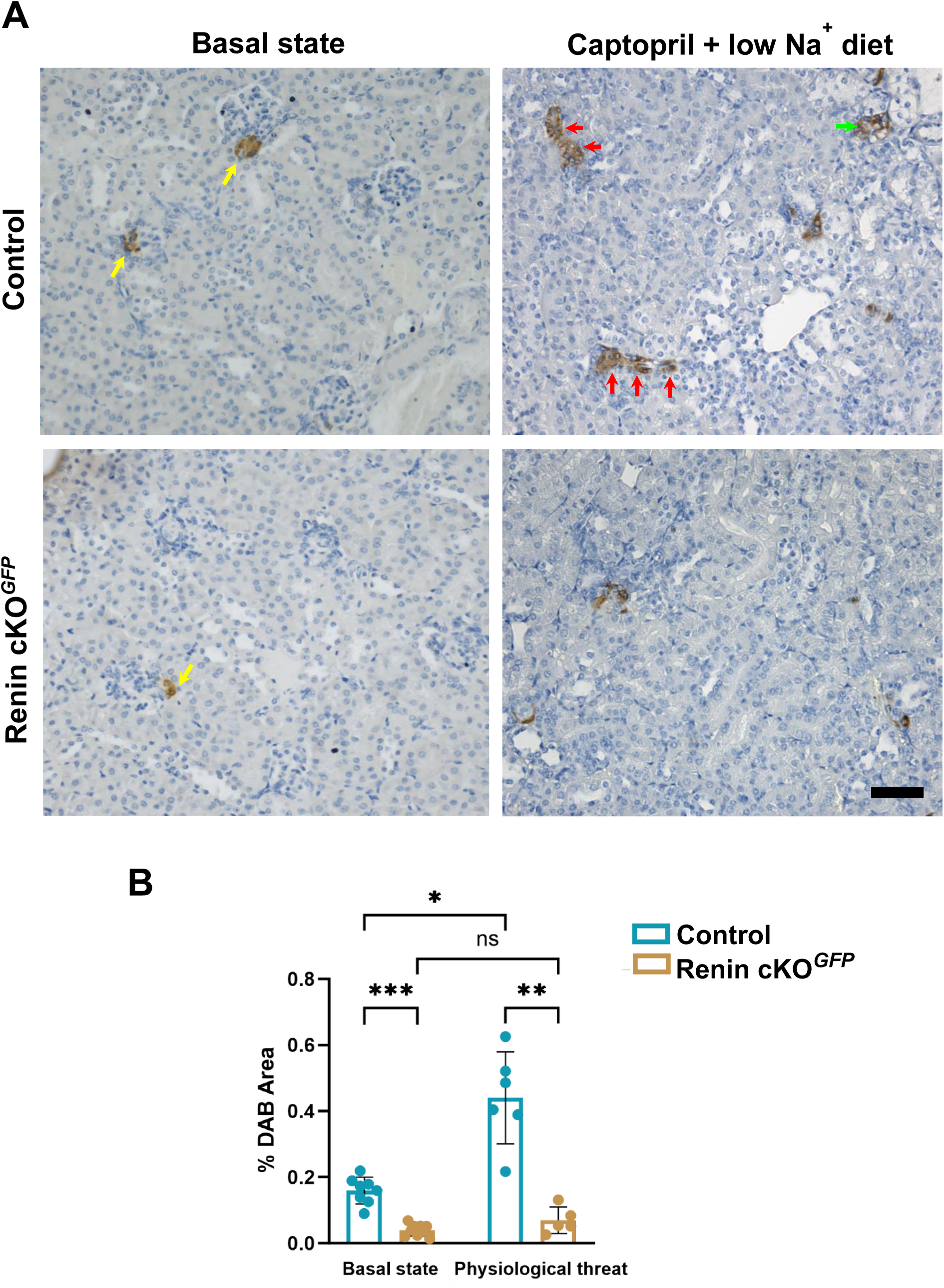
Renin immunostaining of kidney sections from 2-month-old control and *Akr1b7^CreERT2/+^; Ren1^cFl/−^*; R26R*^mTmG^* (Renin cKO*^GFP^*) mice under basal conditions and homeostatic stress. **A.** Results show the normal renin distribution (brown, yellow arrows) at the entrance of the glomeruli in control kidneys under basal conditions whereas a marked decrease in renin distribution and storage is seen in kidneys of Renin cKO*^GFP^* mice. In response to Captopril + low Na^+^ diet treatment, control mice showed renin along the arterioles (red arrows) and in the mesangium (green arrow). The response of Renin cKO*^GFP^* mice was greatly reduced with no visible extension into arterioles and mesangium. Bar: 75 µm. **B.** Quantification of areas of Renin positive immunostaining. We observed a significant decrease in renin immunostaining in mutant mice compared to control animals under basal conditions (**Control-basal: n=8;** Mutant-basal: n=9; ***, *P*<0.001). Similarly, renin immunostaining under recruitment conditions is significantly lower in mutant mice compared to controls, (**Control-recruited: n=6;** Mutant-recruited: n=5; **, *P*<0.01). Under physiological stress, control mice exhibited significantly higher levels of renin expression compared to basal conditions (**Control-basal: n=8;** Control-recruited: n=6; *, *P*<0.05**). Although renin levels in mutant recruited mice were higher than in non-recruited animals the difference was not significant** (Mutant-basal: n=9; Mutant-recruited : n=5; *P*>0.05). Brown-Forsythe and Welch ANOVA followed by Dunnett’s T3 multiple comparisons test.

**Figure 7.**
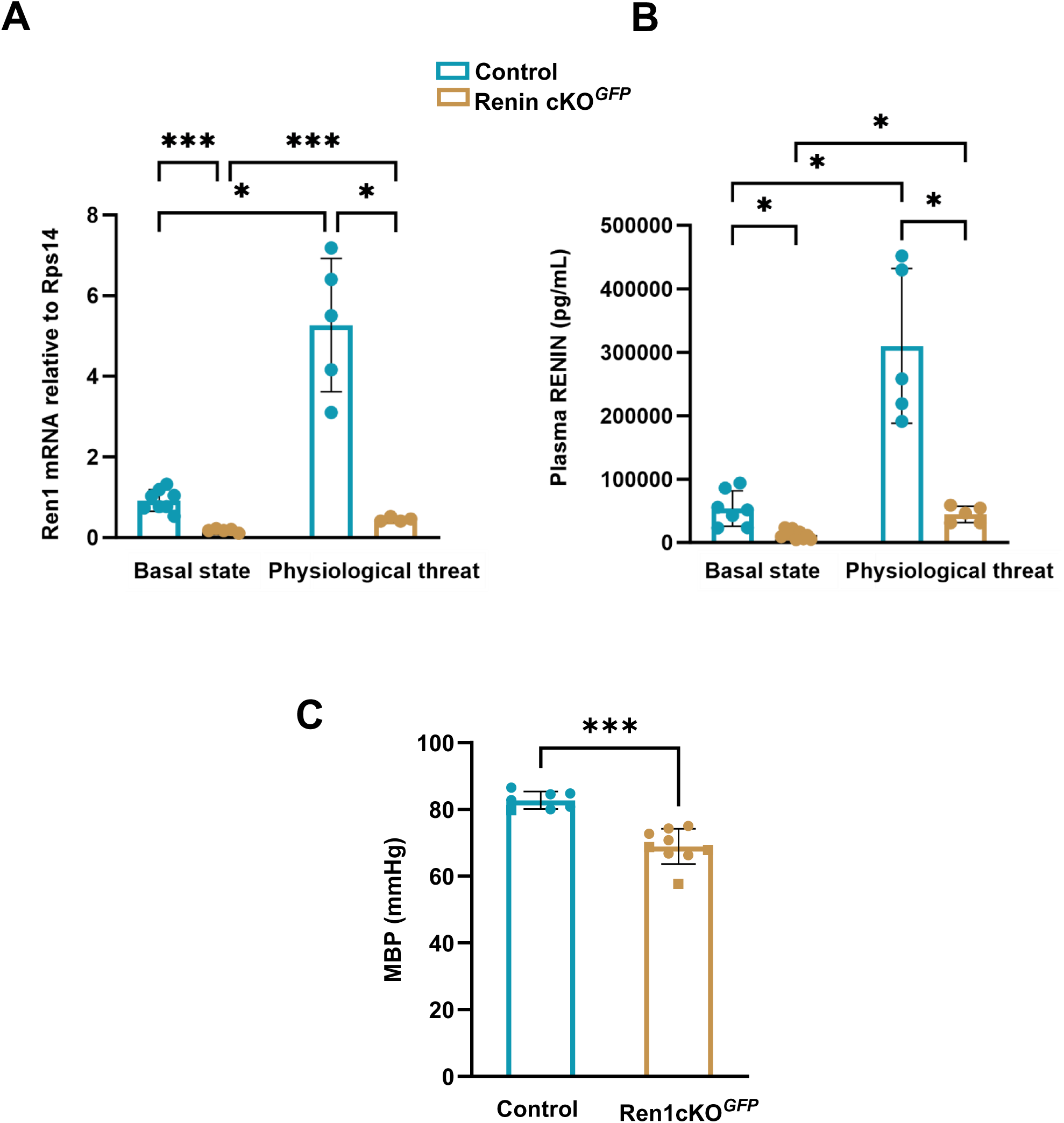
Renin mRNA expression, plasma renin levels and blood pressure in renin Renin cKO mutant mice. Tamoxifen-treated *Akr1b7^CreERt2/+^;Ren1^cFl/−^;R26^mTmG/+^* (Renin cKO*^GFP^*) mice were compared to *Akr1b7^CreERT2/+^; Ren1^c+/−^*; R26R*^mTmG^* control mice under basal conditions and homeostatic stress. Adult (2 to 5-month-old) mice received tamoxifen (2mg/20g BW) i.p. on 5 consecutive days and were analyzed after a washout period of 10–15 days. A group of mice were treated with captopril (0.5 mg/mL) + low Na^+^ (0.1%) diet for the last 8 days of the experiment to induce recruitment of renin cells. **A.** Renin mRNA levels in kidney cortices. We observed a significant decrease in renin mRNA levels in Renin cKO*^GFP^* mutant mice compared to controls in both the basal state (Control-basal: n=8; Mutant basal: n=5; ***, *P*<0.001) and under physiological threat (Control-recruited: n=5; Mutant-recruited: n=4; *8,* *, *P*<0.05). Under physiological stress, control mice exhibited significantly higher levels of renin mRNA compared to basal conditions (Control-basal: n=8; Vontrol-recruited: n=5; *, *P*<0.05). Renin mRNA levels in mutant recruited mice were significantly higher than in non-recruited animals (Mutant-basal: n=5; Mutant-recruited: n=4; ***, *P*<0.001), suggesting that mutant animals are attempting to recruit. **B.** Plasma renin. Circulating renin levels followed a similar trend: Control-basal: n=7 vs Mutant-basal: n=9; *P*<0.05; Control-recruited: n=5 vs Mutant-recruited: n=5; *P*<0.05; Control-basal: n=7 vs Control-recruited: n=5; **\*,** *P*<0.05; Mutant-basal: n=9 vs Mutant-recruited: n=5; *, *P*<0.05. **C**. Arterial blood pressure. Renin mutant mice exhibited a significant decrease in arterial blood pressure under basal conditions compared to controls: Control-basal: n=7 *vs* Mutant-basal: n=9; ***, *P*<0.005. MBP, mean blood pressure. Brown-Forsythe and Welch ANOVA followed by Dunnett’s T3 multiple comparisons test

These results indicate that Renin can be efficiently deleted in the adult using our *Akr1b7^CreERT2^* conditional model.

### Mice develop concentric vascular hypertrophy after *Akr1b7^CreERT2^* mediated renin deletion in the adult

Next, we sought to use the *Akr1b7^CreERT2^* model to determine the effects of long-term deletion of renin in the adult mouse, specifically whether the mice develop concentric vascular hypertrophy upon renin deletion. In these experiments, we administered tamoxifen (2mg/20g BW) i.p. on 5 consecutive days to *Akr1b7^CreERT2/+^;Ren1^cFl/−^;R26^mTmG/+^* (Renin cKO*^GFP^*) and *Akr1b7^CreERT2/+^;Ren1^c+/−^;R26^mTmG/+^* controls at 1 month of age on 5 consecutive days and studied the mice at 5.5 months. Figure 8A shows immunostaining for Acta2 of kidney sections to visualize the vasculature. Renin cKO*^GFP^* kidneys showed thicker arteries and arterioles. This phenotype is similar to the one observed with mutations of the RAS genes or chronic treatment with RAS inhibitors. In addition, renin cells could be tracked by the expression of GFP even when they were unable to express renin (Fig. 8B) allowing the tracking and isolation of these abnormal cells.

**Figure 8.**
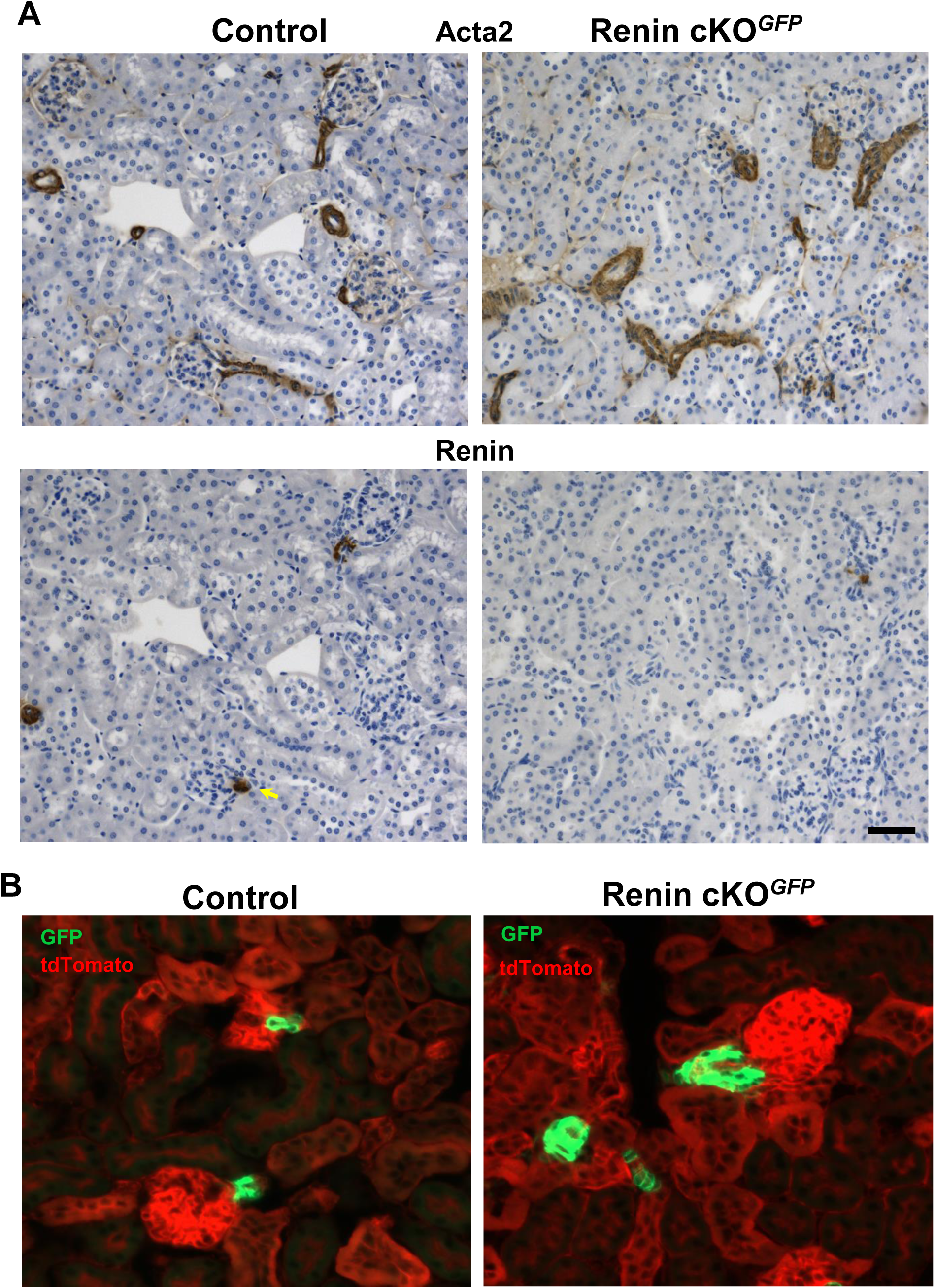
Long term effect of renin deletion in adult animals using the conditional *Akr1b7^CreERt2^* model. Tamoxifen (2mg/20g BW) was administered i.p. on 5 consecutive days *Akr1b7^CreERt2/+^;Ren1^cFl/−^;R26^mTmG/+^* (Renin cKO*^GFP^*) and *Akr1b7^CreERt2/+^;Ren1^c+/−^;R26^mTmG/+^* controls at 1 month of age on 5 consecutive days and the mice were studied at 5.5 months. **A. Top Panel.** Immunostaining for Acta2 of kidney sections from tamoxifen treated Renin cKO*^GFP^* and control mice. Renin cKO*^GFP^* kidneys showed thicker intrarenal arterial and arteriolar (red arrows) walls than controls. This phenotype is similar to the one observed with spontaneous mutations of any of the renin-angiotensin system (RAS) genes or treatment with RAS inhibitors. **Bottom panel**. Renin immunostaining of adjacent sections. Control mice showed the normal renin distribution (brown, yellow arrows) at the entrance of the glomeruli in control kidneys under basal conditions whereas Renin cKO exhibited a marked decrease in renin immunostaining. Bar: 50 µm. **B.** GFP and tdTomato expression in frozen sections of kidneys from the same animals shown in A. Renin cells can be tracked by the expression of GFP even when they are unable to express renin (Fig. 8A) allowing the tracking and isolation of these abnormal cells. Bar: 50 µm.

## DISCUSSION

In this study we describe the first knock-in inducible Cre recombinase mouse model for gene targeting specifically in renin cells. This model consists of a knock-in *P2A-CreERT2* cassette inserted immediately upstream of the TGA stop codon of the mouse *Akr1b7* gene. The *Akr1b7^CreERT2^* model has significant advantages over existing renin cell Cre lines. First, by following a knock-in experimental approach we avoided potential problems associated with transgenic lines that are prone to spurious and/or ectopic gene expression. There is one other reported inducible line where Cre is driven by a *Ren1^c^* promoter as a BAC transgene, and to our knowledge has not yet been fully characterized (16). Second, by using an inducible model we were able to circumvent the confounding developmental effects introduced by non-inducible Cre recombinase lines when the effects of gene deletion are studied in the adult animal. Third, by selecting the independent marker of renin cells, Akr1b7, we ensured that renin expression is not affected by the incorporation of the *CreERT2* cassette at the renin locus. In the kidney, Akr1b7 is expressed exclusively in renin cells. Akr1b7 is co-expressed with renin during development and in the adult under different physiological and pathological conditions (20, 21). Although Akr1b7 and renin are regulated by common factors (21), the expression of one gene does not influence the expression of the other (21, 31). As a result, Akr1b7 can be utilized as a marker for cells that are programmed for the renin phenotype even in the absence of renin (21). Unlike renin, with the exception of the adrenal gland, it is expressed at relatively low levels only in a few organs outside the kidney (32).

The placement of the *P2A-CreERT2* cassette at the Akr1b7 locus had no adverse effects on the mice. Both heterozygous and homozygous *Akr1b7^CreERT2^* mice were viable, fertile, normal in size, and did not display any gross physical or behavioral abnormalities. The function of Akr1b7 remains elusive. Akr1b7 is dispensable for mouse normal development and reproduction (33). Moreover, despite its specific localization in renin cells (20, 21, 31), Akr1b7 does not affect renin expression, localization, and release under normal conditions and in response to physiological stress (31). For this reason, we were unable to evaluate the functional integrity of Akr1b7 in *Akr1b7^CreERT2^* mice. Nevertheless, we observed no difference in the levels and localization of *Akr1b7* mRNA by *in situ* hybridization in the kidney of mice carrying the *Akr1b7^CreERT2^* allele indicating that the genetic manipulation did not affect the expression of endogenous Akr1b7.

Since the *P2A-CreERT2* cassette was inserted upstream of the ATG of the endogenous *Akr1b7* locus, it is expected to be regulated by the same elements that control Akr1b7 expression. Our results show that the fluorescence signal from heterozygous *Akr1b7^CreERT2/+^;R26R^mTmG/+^*and *Akr1b7^CreERT2/+^;R26R^tdTomato/+^* reporter mice clearly aligned with the pattern of endogenous Akr1b7 protein expression during embryonic life, and in the adult under basal conditions and when homeostasis was compromised (20, 21). Notably, within the kidney cortex, *Akr1b7^CreERT2^* mice exhibited Cre recombinase activity specifically in the JG areas. The embryonic and early postnatal *Akr1b7^CreERT2^* expression pattern, encompassing arteries, arterioles, intraglomerular mesangium, and JG areas, underscores the importance of the *Akr1b7^CreERT2^* model as a unique tool to perform inducible gene manipulation in renin cells during development.

In addition to being cell specific, an optimal inducible model must be tightly regulated. Since CreERT2 is constitutively expressed, recombination at LoxP sites might happen even in the absence of tamoxifen. Our results using *Akr1b7^CreERT2/+^*;R26R*^mTmG/+^* mice showed absence of recombination when tamoxifen was not given to the animals, indicating that the system is tightly regulated based on this particular reporter. In addition, in adult animals treated with tamoxifen, the GFP signal was confined strictly to JG cells with no expression in other portions of the vasculature which would have resembled the embryonic pattern, indicating that recombination did not occur before induction with tamoxifen.

Akr1b7 is expressed in a few organs outside the kidney, and in most cases at very low levels (32). However, Akr1b7 is highly expressed in the murine adrenal gland where it is confined to the zona fasciculata of the cortex (28, 33, 34). The cells in this area of the adrenal cortex synthesize glucocorticoids and androgens, and it has been suggested that Akr1b7 may be involved in the detoxification of harmful aldehydes produced during the synthesis of steroids (34). In the adrenal gland of tamoxifen-induced *Akr1b7^CreERT2/+^;R26R^mTmG/+^* reporter mice, we found high levels of GFP expression exclusively in the zona fasciculata of the cortex. This finding suggests that the *Akr1b7^CreERT2^* model may be a valuable resource for lineage tracing and gene targeting in the glucocorticoid and androgen-producing cells of the adrenal gland. Also, it seems that no obvious defects in the adrenal were found.

We used double reporter mice, *Akr1b7^CreERT2^;Ren1^cYFP^;R26R^tdTomato^* to evaluate the degree of co-localization of renin and Akr1b7^CreERT2^ as well as the rate of recombination after tamoxifen induction. The expression of YFP driven by the *Ren1^c-YFP^* transgene during development, in adult life, and in response to physiological stress (i.e. recruitment) precisely follows the pattern of renin expression visualized by immunostaining (23). Our results show that all YFP (renin) expressing cells were positive for tdTomato (*Akr1b7^CreERT2^*) strongly suggesting that *Akr1b7^CreERT2^* confers not only renin cell-specific *Cre* activity but also a high rate of recombination after tamoxifen induction. This conclusion is also supported by the lack of significant difference between the JGAi determined using YFP^+^ cells and tdTomato^+^ cells. Our observation that both reporters were highly co-localization in JG cells, along the arterioles and the intraglomerular mesangium under conditions of physiological stress suggests that *Akr1b7^CreERT2^* model can also be used for the genetic manipulation of cells during reacquisition of the renin phenotype.

The activation of reporter alleles using Cre-LoxP systems does not indicate that the recombination necessarily will take place at other floxed loci in the cell, a process known as non-parallel recombination (35). It has been reported that the rate of Cre mediated recombination depends on several factors, including the distance between the LoxP sites, and the chromosomal location and epigenetic context of the floxed alleles (34 and references therein). In order to investigate the potential use of the *Akr1b7^CreERT2^* model for genetic manipulation of genes beyond well-established reporter floxed alleles, we generated *Akr1b7^CreERT2/+^;Ren1^cFl/+^;R26^mTmG/+^* mice to target renin in the adult animal. Compared to controls, renin mutant animals exhibited significantly lower expression of renin in the kidney at both mRNA and protein levels, with only a few renin positive cells detectable by immunostaining. In addition, mutant mice exhibited significantly lower levels of circulating renin and low blood pressure. These results demonstrate that renin can be efficiently deleted in the adult using our *Akr1b7^CreERT2^* conditional model. Furthermore, the response of renin mutant mice to a physiological stress, i.e. captopril + low Na^+^ diet, was severely compromised. When homeostasis is threatened (e.g. dehydration, hemorrhage, hypoxemia, and deletion of RAS genes), Akr1b7 is upregulated, along with renin, in recruited arteriolar smooth muscle cells, mesangial cells, and interstitial cells (21), The lack of renin expression in these cells is probably the result of further renin deletion as Akr1b7 gets activated during recruitment in renin mutant mice.

Our group and others have shown that experimental or spontaneous mutations of any of the RAS genes or long-term treatment with inhibitors of the RAS in mammals, including humans, lead to the development of concentric arterial and arteriole hypertrophy (7–15). We found that mice with ablation of renin cells using Diphtheria toxin (36) or conditional deletion of Integrin β1 (15) do not develop kidney vascular hypertrophy indicating that renin cells *per se* are responsible for the vessel thickening. Furthermore, we demonstrated that vascular hypertrophy in renin-deficient mice is not the result of the proliferation of renin cells (37) but instead is due to their transformation from an endocrine to an invasive matrix-secretory phenotype accompanied by inward accumulation of smooth muscle cells (15). These observations indicate that renin cells are not proliferative and they increase their numbers by phenotypic switching and cell transformation.

Nevertheless, the underlying mechanisms leading to this vascular pathology remain unknown. In this study, we show that deletion of renin during adulthood leads to the development of concentric vascular hypertrophy resulting in a marked reduction of the vessels lumen. This morphological phenotype is similar to the one observed in animals with deletion of renin globally or constitutively in renin cells (9, 13). However, the latter show additional renal abnormalities, including papillary atrophy, underdeveloped medulla, hydronephrosis, interstitial fibrosis, focal glomerulosclerosis, and perivascular infiltration of mononuclear cells. These morphological alterations can be the result of developmental changes caused by the absence of renin expression throughout the life of the animals and may alter the vascular phenotype. Our inducible model offers the possibility to study the vascular hypertrophy independently of these confounding factors. In addition, our model allows the labeling of the abnormal vascular cells for isolation and exploring the molecular pathways and cellular interactions underlying the observed vascular hypertrophy. Further studies with these animals will help elucidate the specific role of renin deficiency in the development of concentric vascular hypertrophy. Additionally, unraveling the signaling pathways triggered by renin deficiency may uncover novel therapeutic targets for conditions associated with vascular hypertrophy.

In conclusion, the *Akr1b7^CreERT2^* mouse showed high levels of tamoxifen-induced recombination that is renin cell-specific and tightly regulated. The *Akr1b7^CreERT2^* model constitutes an excellent tool for the temporal and spatial control of target genes and for lineage tracing during development and in adult mice.

## ACKNOWLEDGEMENTS

We thank Xiuyin Liang, Fang Xu, Minghong Li, Thomas Wagamon and DJ White for excellent technical assistance. This work used ZEISS Lightsheet7 and Imaris software in the Advanced Microscopy Facility which is supported by the University of Virginia School of Medicine, Research Resource Identifiers (RRID): SCR_018736.

