## Supplementary Figures for "An efficient inducible model for the control of gene expression in renin cells"

**Supplementary Figure 1. Expression of *Akr1b7* in the kidney cortex of *Akr1b7^CreERT2^* Supplementary Figure 1. Expression of *Akr1b7* in the kidney cortex of *Akr1b7^CreERT2^* mice.** Shown are images of *in situ* hybridization for *Akr1b7* mRNA. **A.** Mouse wild-type for the *Akr1b7* allele treated with three consecutive i.p injections of corn oil (0.1 cc/20 g) and analyzed 3 weeks after the last injection. **B.** Mouse heterozygous for the *Akr1b7^CreERT2^* allele treated with three consecutive i.p injections of corn oil (0.1 cc/20 g) and analyzed 3 weeks after the last injection. **C.** Mouse heterozygous for the *Akr1b7^CreERT2^* allele treated with three consecutive i.p injections of tamoxifen (2 mg/20 g) and analyzed 3 weeks after the last injection. Left panels show multiple JG areas positively stained for *Akr1b7* in all conditions. *Akr1b7* staining was restricted to the JG cells. Right panels show higher magnification images of *Akr1b7* positive JG cells. Three mice in each group were analyzed. We observed no difference in the levels and localization of *Akr1b7* mRNA in the kidney of mice carrying the *Akr1b7^CreERT2^* allele. In addition, *Akr1b7* mRNA expression was not affected by tamoxifen administration. Bars: low magnification: 500 µm; high magnification: 100 µm.

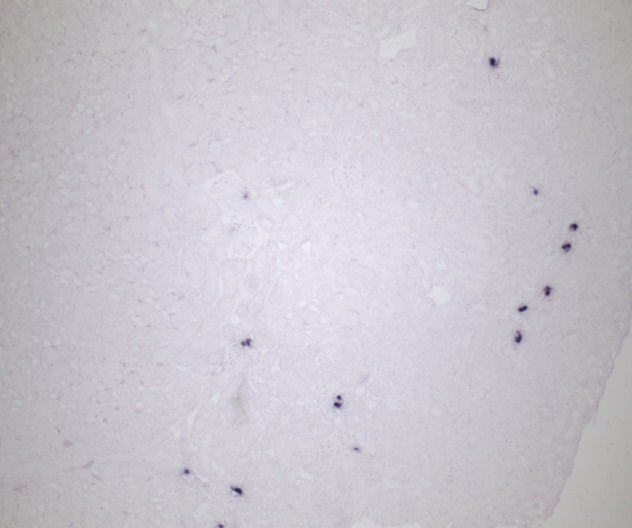

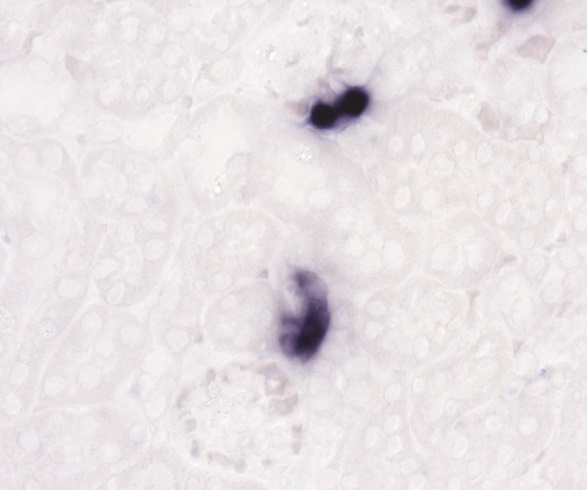

**A**

***Akr1b7*^+/+^ Vehicle**

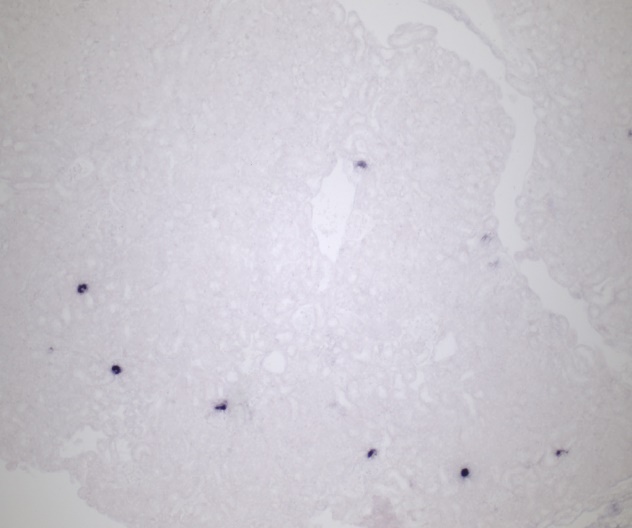

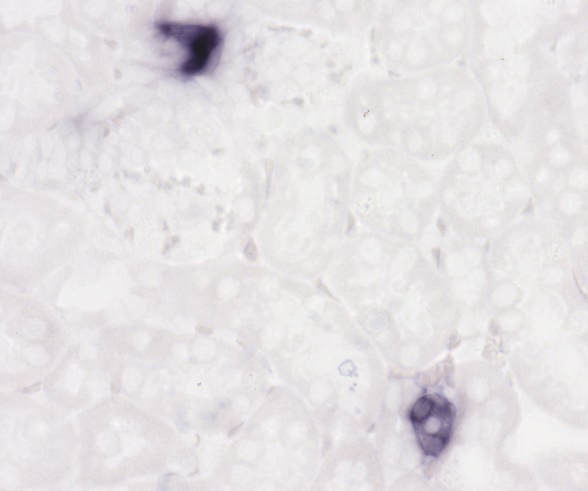

***Akr1b7*^CreERT2/+^ Vehicle**

**B**

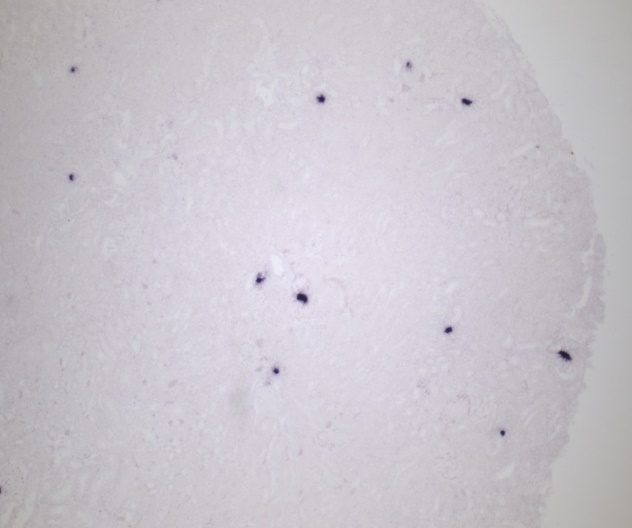

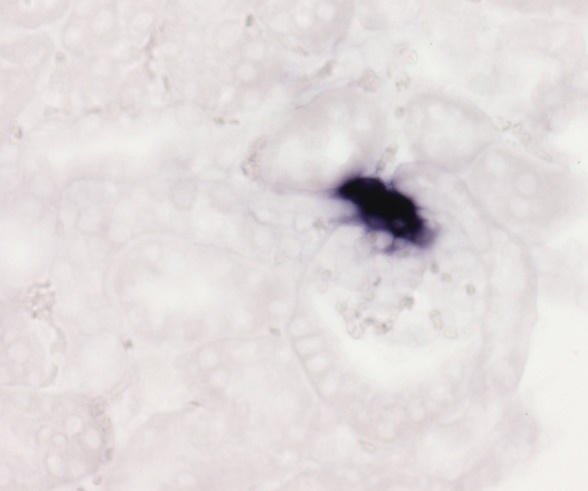

***Akr1b7*^CreERT2/+^ Tamoxifen**

**C**

**Adrenal Gland**

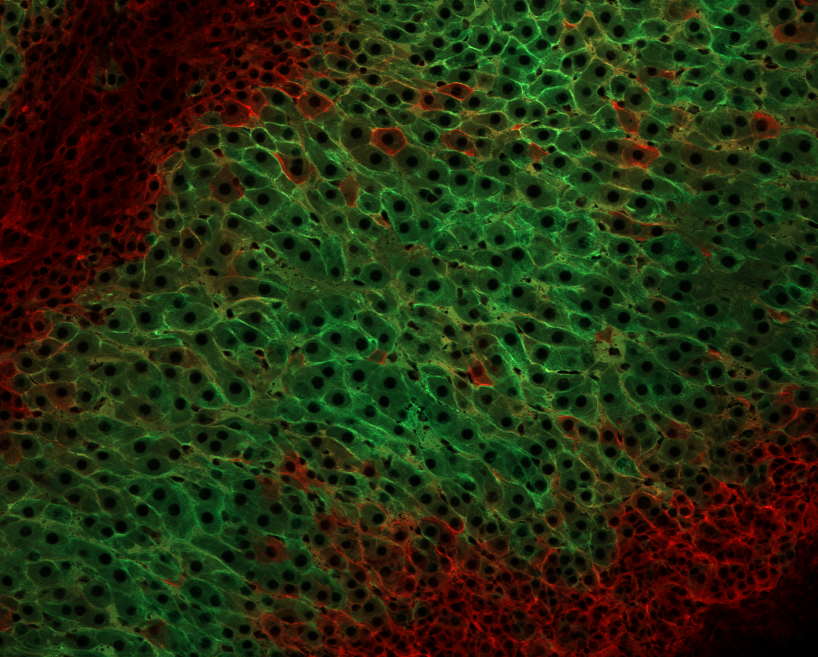

**M**

**ZF/ZR**

**ZG**

**GFP**

**tdTomato**

**Supplementary Figure 2. Pattern of Cre expression in the adrenal glands** **of *Akr1b7^CreERT2^* reporter mice.** GFP and tdTomato expression in frozen sections of adrenal glands from tamoxifen-treated Akr1b7*^CreERT2^*;R26R*^mTmG^* mice at P30. We observed intense GFP signal in cells in the zona fasciculate of the adrenal cortex as previously described in mice and humans (28). M, medulla; ZG, zona glomerulosa; ZF, zona fasciculata, ZR; zona reticularis, Bar: 50 µm.
