## Supplementary Video Legends for "An efficient inducible model for the control of gene expression in renin cells"

**Video 1.** **Reconstructed 3D image of kidney Akr1b7*^CreERT2^*;R26R*^tdTomato^* mice under normal condition, related to Figure 3**

Red fluorescence indicates Akr1b7*^CreERT2^* expression upon tamoxifen administration and GPF shows immunofluoresnce for ACTA2. Under basal conditions, tdTomato signal was clearly localized to the tip of the arterioles in the JG areas.

**Video 2. Reconstructed 3D image of kidney Akr1b7*^CreERT2^*;R26R*^tdTomato^* mice under physiological stress, related to Figure 3**

Red fluorescence indicates Akr1b7*^CreERT2^* expression upon tamoxifen administration and GPF shows immunofluoresnce for ACTA2. Under physiological stress, tdTomato signal was much brighter, and the labelled JG areas looked larger compared to controls. In addition, the tdTomato signal extended into the arterioles as expected in recruited mice.
